# WEE1 inhibition delays resistance to CDK4/6 inhibitor and antiestrogen treatment in estrogen receptor-positive breast cancer

**DOI:** 10.1101/2024.09.15.613122

**Authors:** Wei He, Diane M. Demas, Pavel Kraikivski, Ayesha N. Shajahan-Haq, William T. Baumann

## Abstract

Although endocrine therapies and Cdk4/6 inhibitors have produced significantly improved outcomes for patients with estrogen receptor positive (ER+) breast cancer, continuous application of these drugs often results in resistance. We hypothesized that cancer cells acquiring drug resistance might increase their dependency on negative regulators of the cell cycle. Therefore, we investigated the effect of inhibiting WEE1 on delaying the development of resistance to palbociclib and fulvestrant. We treated ER+ MCF7 breast cancer cells with palbociclib alternating with a combination of fulvestrant and a WEE1 inhibitor AZD1775 for 12 months. We found that the alternating treatment prevented the development of drug resistance to palbociclib and fulvestrant compared to monotherapies. Furthermore, we developed a mathematical model that can simulate cell proliferation under monotherapy, combination or alternating drug treatments. Finally, we showed that the mathematical model can be used to minimize the number of fulvestrant plus AZD1775 treatment periods while maintaining its efficacy.

## Introduction

Breast cancer is a commonly diagnosed cancer among women, with approximately 310,720 new cases expected to be diagnosed in 2024 in the United States^1^. Despite advances in treatment options, it is estimated that 42,250 women will die as a result of invasive breast cancer in 2024^1^. Targeted therapies play an important role in the treatment of estrogen receptor-positive (ER+) breast cancer, which is the most common subtype of breast cancer^2^. Endocrine therapies, the cornerstone of targeted treatment for ER+ breast cancer, interfere with estrogen signaling, which promotes the growth and proliferation of cancer cells^3^. Antiestrogens have revolutionized the treatment of breast cancer, resulting in dramatic improvements in long-term survival rates while avoiding the toxicity of traditional chemotherapies. In addition to endocrine therapy, cyclin-dependent kinase (Cdk) 4 and 6 inhibitors, such as palbociclib, ribociclib and abemaciclib, have become a significant targeted therapy for the treatment of ER+ breast cancer^4^. These inhibitors halt cancer progression by blocking the activity of Cdk4/6, a critical complex in cell cycle progression and cell proliferation^4^. Cdk4/6 inhibitors in combination with endocrine therapy are the standard-of care treatment for most patients with ER+ advanced breast cancer^4^.

Although most cancer cells are responsive to the targeted therapies, continuous application of targeted drugs often results in acquired resistance, which remains one of the major impediments to treating cancer^5,6^. Without exception, acquired resistance to endocrine therapy and Cdk4/6 inhibitors inevitably ensues^4,7–11^. Cancer cells’ plasticity can drive their transformation towards a state where they can temporarily adapt to the presence of the targeted drug, leading to reduced drug sensitivity^12,13^. This acquired resistance can potentially be reversed by discontinuing the drug, to allow sensitivity to return, or it may become a permanent state and drive relapse^5^. The mechanisms of reversible drug resistance are dynamic and potentially can be overcome by modifying treatment strategies or removing the pressure exerted by the targeted drug. Overcoming reversible drug resistance is a challenging problem, which often involves using a new combination or alternating therapy. By targeting multiple pathways involved in cell growth and survival, as well as vulnerabilities within cancer cells, combination treatments can reduce the possibility of resistance development^14,15^. Alternating treatments sequentially administer different drugs over time, regularly changing the selective pressure exerted on cancer cells to avoid selecting for resistance mechanisms and making it more difficult for cancer cells to adapt and develop resistance to single drug^16–18^.

WEE1 is a protein of the tyrosine kinase family that regulates the G2 checkpoint^19^. WEE1 inhibits Cdk1 by phosphorylating Tyr15 (Y15), stopping cells from entering mitosis to allow time for DNA repair^20^. While WEE1 kinase primarily functions at the G2/M checkpoint in the cell cycle, it can also have regulatory roles in the G1/S transition and S phase^21–27^. When cells reach S phase, replication is initiated from a large number of replication origins and the absence of WEE1 leads to a Cdk-dependent increase in replication initiation^23^. The lack of Cdk inhibition also leads to unscheduled initiation of replication. Loss of WEE1 can lead to unscheduled, excessive origin firing resulting in exhaustion of nucleotides^23^ and replication catastrophe^28^. The deficiency of nucleotides triggers replication fork slowing and stalling, resulting in accumulation of single-stranded DNA (ssDNA) at the stalled forks –– a situation called replication stress (RS). RS in cancer cells can be induced by oncogenic activities and aggravated in resistant cells^29–31^. A therapeutic strategy that exacerbates the existing RS in cancer cells can drive cancer-specific cell death^32,33^. Moreover, it is reported that elevated CyclinE1 will impair DNA replication causing replication stress^34,35^. Single cell sequencing revealed that overexpression of CyclinE1 results in replication stress and WEE1 inhibition exacerbated the stress induced by CyclinE1 overexpression and caused cytotoxicity. Downregulation of CyclinE1 rescued the increased sensitivity to WEE1 inhibitors^36^. CyclinE1 expression is also associated with high levels of replication stress in triple-negative breast cancer^37^. Upregulation of Cdk6 to overcome epidermal growth factor receptor (EGFR) inhibitor-induced G1/S phase cell cycle arrest is also associated with increased RS^38^. In contrast, a decrease in G1 and S phase Cdk activity can confer resistance to WEE1 inhibition, providing evidence that major cytotoxic effects of WEE1 inhibition are exerted in S phase^23,39^.

At the same time, CyclinE1 and Cdk6 up-regulation are two of the many mechanisms claimed to contribute to the acquired resistance to the Cdk4/6 inhibitor palbociclib and the endocrine therapy fulvestrant (faslodex; ICI 182,780; ICI), respectively^40–46^. We hypothesized that the ER+ breast cancer cells developing resistance to palbociclib or ICI will exhibit increased G1 and S phase Cdk activity, which will exacerbate the existing RS and sensitize them to WEE1 kinase inhibition^47–49^. Then, WEE1 inhibitors could particularly target those cells using a relatively low dose. Therefore, we tested if alternating palbociclib and a combination of ICI plus a WEE1 small molecule inhibitor, AZD1775 (Adavosertib), would delay or preclude the development of resistance to palbociclib and ICI. While AZD1775 has shown promising antitumor activity, a major concern regarding AZD1775 is that it is not well tolerated by patients and has been associated with certain toxicities, subjecting the drug to criticism and scrutiny^50,51^. To limit the toxicity of AZD1775, we fixed its dose at a low concentration of 250nM in our alternating treatment. Also, alternating treatment by itself has been shown to reduce the toxicity of a WEE1 inhibitor.

To better understand the possible interactions among these three drugs and to be able to make predictions about future therapy options, we built a mathematical model to simulate cell proliferation under monotherapy, combination and alternating treatments. We used increased CyclinE1 and Cdk6 protein level changes under palbociclib and ICI treatments as the resistance mechanisms causing the resumption of proliferation under the monotherapy treatments. This resumption of proliferation can be reduced or eliminated by adding AZD1775 into the mix. More importantly, to relieve the potential side effects and toxicity of AZD1775, we showed how the mathematical model can be used to minimize the number of ICI plus AZD1775 periods in the alternating treatment while still maintaining the efficacy of AZD1775.

## Results

### Alternating treatment of palbociclib with ICI 182,780 plus AZD1775 delays the development of resistance compared to monotherapy and alternating therapy without AZD1775

Different 12-month drug treatment strategies for MCF7 breast cancer cells are shown in Fig. 1. MCF7 cells are treated either by palbociclib monotherapy, ICI monotherapy, palbociclib alternating with ICI (Alter 1) or palbociclib alternating with ICI plus AZD1775 (Alter 2) over a 12-month period. The alternation interval is four weeks and MCF7 cells are re-plated at the beginning of each month for all treatments. Fig. 2a plots the cell proliferation results with the different drug treatments over 12 months. For comparison, it should be noted that without drug treatment, MCF7 cells will expand over 120-times in 11 days (Supplementary Fig. 1A).

**Fig. 1.**
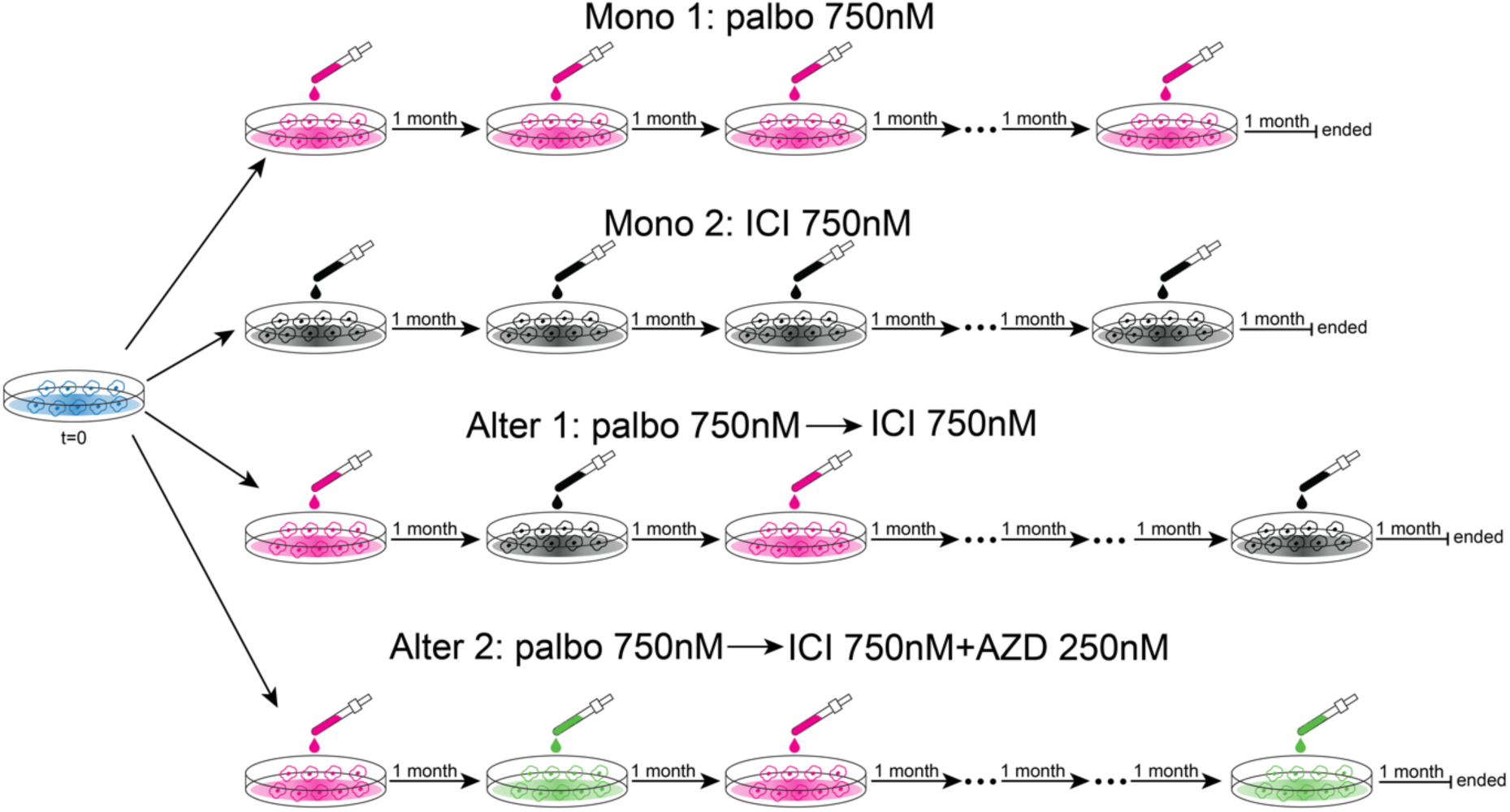
Schematic representation of four 12-month treatment strategies. MCF7 cells are treated by monotherapy or alternating treatment over 12 months, 28 days per month. MCF7 cells are re-plated at the beginning of each month in the mono and alternating treatments. Mono 1: palbociclib monotherapy at 750nM. Mono 2: ICI monotherapy at 750nM. Alter 1: alternating treatment of palbociclib 750nM with ICI 750nM, drug used is altered at the beginning of each month. Alter 2: alternating treatment of palbociclib 750nM with ICI 750nM plus AZD1775 250nM, drug used is altered at the beginning of each month. Palbo: palbociclib, AZD: AZD1775.

**Fig. 2.**
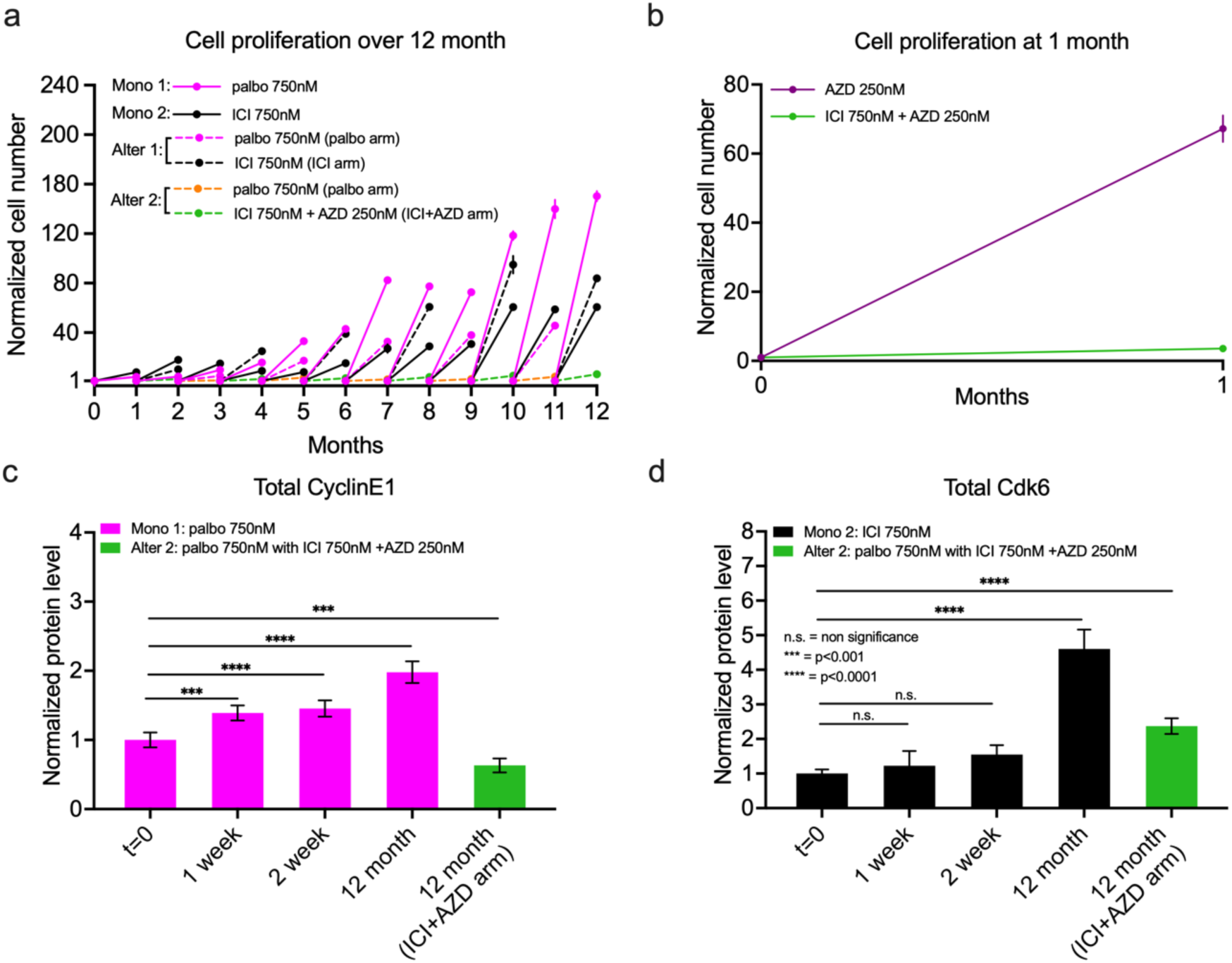
Cell proliferation and protein level changes after 12 months mono and alternating treatments. **a** Experimental cell proliferation data (mean ± s.d., n = 3, technical replications) for palbociclib monotherapy (solid magenta line), ICI monotherapy (solid black line), alternating treatment of palbociclib (dashed magenta line) with ICI (dashed black line), and alternating treatment of palbociclib (dashed orange line) with ICI plus AZD1775 (dashed green line). Experimental settings are the same as Fig. 1. The cell number is normalized to the cell number at re-plating. **b** Experimental cell proliferation data (mean ± s.d., n = 3) for AZD (solid purple line) and ICI plus AZD1775 (solid green line) at 1 month (28 days). **c** Bar plot of western blot data for total CyclinE1 (mean ± s.d., n = 3) level changes in palbociclib monotherapy and alternating treatment of palbociclib with ICI plus AZD1775 at different time points. Statistical testing was performed by one-way ANOVA. Only the significant differences between treatment timepoints and t=0 are shown. **d** Bar plot of western blot data for total Cdk6 (mean ± s.d., n = 3) level changes in ICI monotherapy and alternating treatment of palbociclib with ICI plus AZD1775 at different time points. Statistical testing was performed by one-way ANOVA. Only the significant differences between treatment timepoints and the t=0 are shown. Palbo: palbociclib, AZD: AZD1775.

From Fig. 2a it is seen that palbociclib treatment at 750nM significantly inhibited the cell proliferation during the first 3 months, only allowing the MCF7 cells to proliferate less than 10-fold per month. However, MCF7 cells start to show significant proliferation after 4 months of palbociclib treatment and become increasingly resistant to the drug under continuing treatment. This acquired resistance to palbociclib is fully developed after 10 months with the cells proliferating about 140-fold (full confluence) per month. As expected, the situation for ICI monotherapy is similar, where ICI can effectively inhibit cell proliferation when initially introduced but then it loses its effectiveness over time. While proliferating 8-fold per month initially, MCF7 cells developed the ability to proliferate 60-fold per month after 12-months of ICI monotherapy.

The resistance that develops as the cells adapt to a constant selective pressure necessitates the consideration of innovative combination and sequential treatment strategies. We first tested whether a simple alternation of palbociclib with ICI could affect the development of resistance to both drugs. In Fig. 2a, it is seen that alternating palbociclib with ICI slows down the development of resistance to palbociclib but not ICI. At month 11, comparing the proliferation between the palbociclib monotherapy and the palbociclib treatment arm in the alternating therapy, the cell proliferation is lower in the alternating treatment than in the monotherapy. This difference may be due to the alternation with ICI or may simply be due to the total palbociclib treatment time, which is less in the alternating treatment than in the monotherapy at month 11. In the ICI case, we see that at month 12, although the total ICI treatment time in the alternating therapy is shorter than in the ICI monotherapy, the cell proliferation is higher in the alternating treatment than in the ICI monotherapy. Overall, merely alternating palbociclib with ICI does not effectively suppress cancer cell proliferation over the long term. The breast cancer cells show enhanced proliferation at the end of the treatment and could even develop cross resistance to both of the drugs^55^.

Next, based on our hypothesis, we tested the inhibition effect of alternating palbociclib treatment with ICI plus AZD1775. As shown in Fig. 2a, this alternating treatment successfully inhibited MCF7 cell proliferation over 12 months. The proliferation rate is nearly constant from the beginning to the end of the treatment. Therefore, this alternating protocol can delay the development of resistance to palbociclib and ICI for at least 12 months. To show that the inhibition under ICI plus AZD1775 treatment is not solely from AZD1775, we plotted the cell fold-change in 1 month under AZD1775 treatment and ICI plus AZD1775 treatment in Fig. 1b. We can see that the cell population increased over 60 times in 1 month under 250nM AZD1775 treatment and about 4 times under ICI+AZD1775 treatment. Thus, the combination of ICI and AZD1775 is necessary in the alternating treatment. It is possible that ICI and AZD1775 are targeting different subpopulations of the MCF7 cells during the alternating treatment.

### ICI 182,780 plus AZD1775 in the alternating treatment mitigates the upregulation of Cdk6 in ICI 182,780 treatment and CyclinE1 in palbociclib treatment

CyclinE1 up-regulation is one of the mechanisms claimed to contribute to acquiring resistance to palbociclib^40,43^. Also, in clinical studies, patients with breast cancer who exhibit upregulation of CyclinE1 have poorer responses to palbociclib treatment^41,42^. We measured CyclinE1 level changes by western blot at different timepoints during the palbociclib monotherapy and the palbociclib and ICI plus AZD1775 alternating treatment. In Fig. 2c, the CyclinE1 level gradually increases in response to 12-month palbociclib monotherapy, as would be expected for a long-term resistance mechanism to palbociclib. We can also see that the normalized CyclinE1 level is lower than 1 in the ICI plus AZD1775 arm after 12 months of the alternating treatment. This effect could be due to a combination of ICI treatment and AZD1775 treatment^7,47–49^.

Similarly, Cdk6 overexpression is one of the mechanisms reported to contribute to acquiring resistance to ICI. Increased expression of Cdk6 has been seen in ICI resistant cells^44–46^. Also, breast cancer patients with high Cdk6 levels show significantly shorter progression-free survival on fulvestrant treatment^44^. We measured Cdk6 level changes by western blot at different timepoints during the ICI monotherapy. In Fig. 1d, the Cdk6 level steadily increases in response to long-term ICI treatment. However, ICI plus AZD1775 in the alternating treatment mitigates the upregulation of Cdk6, providing a possible mechanism for how AZD1775 thwarts ICI resistance.

### MCF7 cells exhibited varying drug dose responses following different treatment strategies

At the end of 12 months, a 6-day palbociclib dose-response assay was used to compare the proliferation of MCF7 cells after undergoing no treatment, monotherapy or one of the two different alternating treatments. Fig. 3a-c show the results for three normalizations: growth in vehicle, initial cell numbers at t=0, and the growth-rate inhibition metric, GR^56^. Fig. 3a normalizes the cell proliferation under each condition to its proliferation in vehicle. The parental MCF7 cells are the most sensitive to palbociclib. Compared with the palbociclib monotherapy, MCF7 cells subjected to the two alternating treatment regimens are more sensitive to palbociclib, with no significant difference between the two. When the dose-response results are normalized to t = 0, as shown in Fig. 3b, MCF7 cells exposed to the different treatment conditions exhibit reduced proliferation rates compared to the untreated case. These metrics of drug dose response can be significantly influenced by the number of cell divisions, as has been noticed previously and drove the development of the new GR metric. GR is robust to variations in cell growth rate and quantifies the efficacy of a drug on a per-division basis, which ensures that fast– and slow-dividing cells responding equally to a drug are scored equivalently^56^. Fig. 3c shows the GR values representing the palbociclib dose response for MCF7 cells subjected to the different treatments. In addition to the significant difference between the monotherapy and alternating treatments, the cells undergoing the alternating treatment including AZD1775 are more sensitive to palbociclib compared with the cells undergoing the alternating treatment without AZD1775. Furthermore, no significant difference in response was observed between the parental cells and the cells subjected to 12 months of alternating treatment using palbociclib and ICI plus AZD1775. This suggests that alternating treatment not only can delay the development of resistance to palbociclib but also maintain a sensitivity to palbociclib after 12-months of treatment akin to parental cells, if the alternating drug combinations are selected carefully.

**Fig. 3.**
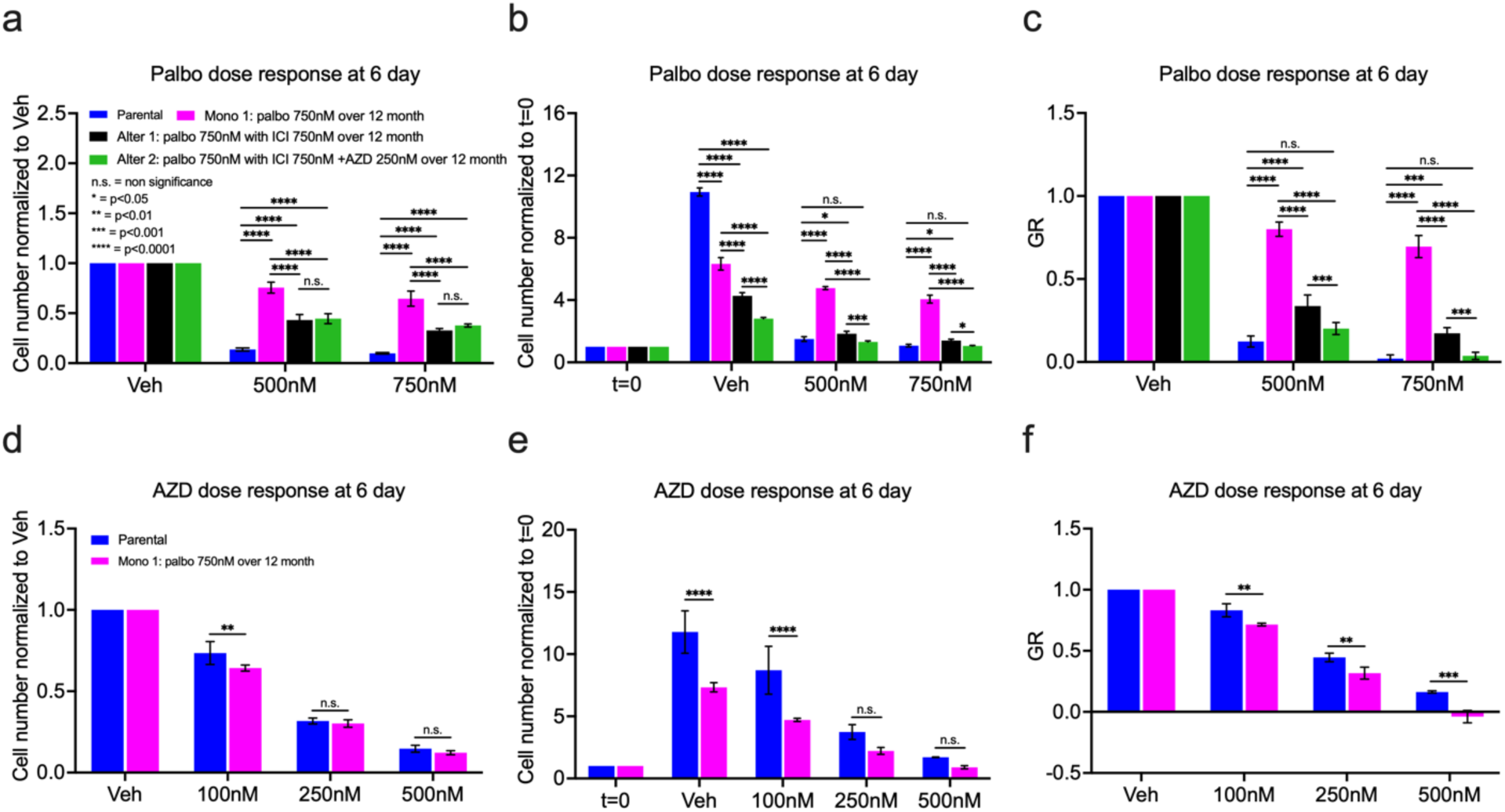
Palbociclib and AZD1775 dose response for MCF7 cells under mono and alternating treatments. **a** Palbociclib dose response normalized to vehicle after 12 months palbociclib monotherapy, alternating treatment of palbociclib with ICI, alternating treatment of palbociclib with ICI plus AZD1775 compared to parental cells. The palbociclib monotherapy and alternating treatments are the same as Fig. 1. **b** Palbociclib dose response normalized to t=0, otherwise same as **a**. **c** The GR value of palbociclib dose response, otherwise same as **a**. **d** AZD1775 dose response normalized to vehicle after 12 months palbociclib monotherapy compared to parental cells. The palbociclib monotherapy is the same as Fig. 1. **e** AZD1775 dose response normalized to t=0, otherwise same as **d**. **f** The GR value of AZD1775 dose response, otherwise same as **d**. Palbo: palbociclib, AZD: AZD1775.

To explore the sensitivity of palbociclib-resistant MCF7 cells to AZD1775, a dose-response experiment using AZD1775 was conducted on the MCF7 cells that underwent 12 months of palbociclib monotherapy. The results are shown in Figs. 3d-f. In particular, the GR values demonstrate that the palbociclib-resistant MCF7 cells exhibit significant sensitivity to AZD1775 at all tested doses. Moreover, at 500nM, AZD1775 induced cytotoxic effects in palbociclib-resistant MCF7 cells, as evidenced by the negative GR value^56^, whereas no such effect was observed in parental MCF7 cells. This result aligns with the idea that CyclinE1 overexpression in breast cancer can lead to Cdk2-dependent replication stress, rendering the cells more sensitive to AZD1775^48^.

### The proposed mathematical model can recapitulate the experimental proliferation data

Systematic application of mathematical models can enhance our understanding of cancer treatment dynamics and can be used to propose optimized treatment regimens that maximize therapeutic efficacy while limiting side effects. The quantities being optimized include drug dosing schedules, sequencing of treatments, and combinations of therapies^57^. We built an ordinary differential equations (ODEs) mathematical model utilizing established mechanisms from the literature to describe the changes in protein expression and proliferation of MCF7 cells in response to various treatment strategies over a period of 12 months^52,53^. The mathematical model is based on the effects of estrogen signaling and Cdk4/6 inhibition on the principal interactions of the G1-S transition, Fig. 4. To capture the progressively increased cell proliferation under palbociclib and ICI treatment, we employed upregulation of CyclinE1 in response to palbociclib and upregulation of Cdk6 in response to ICI. Out of the numerous mechanisms associated with palbociclib and ICI resistance, we selected CyclinE1 and Cdk6 due to their presence in our modeled pathways and experimental data. We calibrated the model to recapitulate MCF7 cell proliferation under the various treatments and showed it is possible for the cells to acquire resistance to the drugs over time by upregulation of the two protein levels. We demonstrated how the effect of AZD1775 decreasing CyclinE1 and Cdk6 levels can overcome the emerging resistance and allow control of cell proliferation in the alternating treatment over 12 months.

**Fig. 4.**
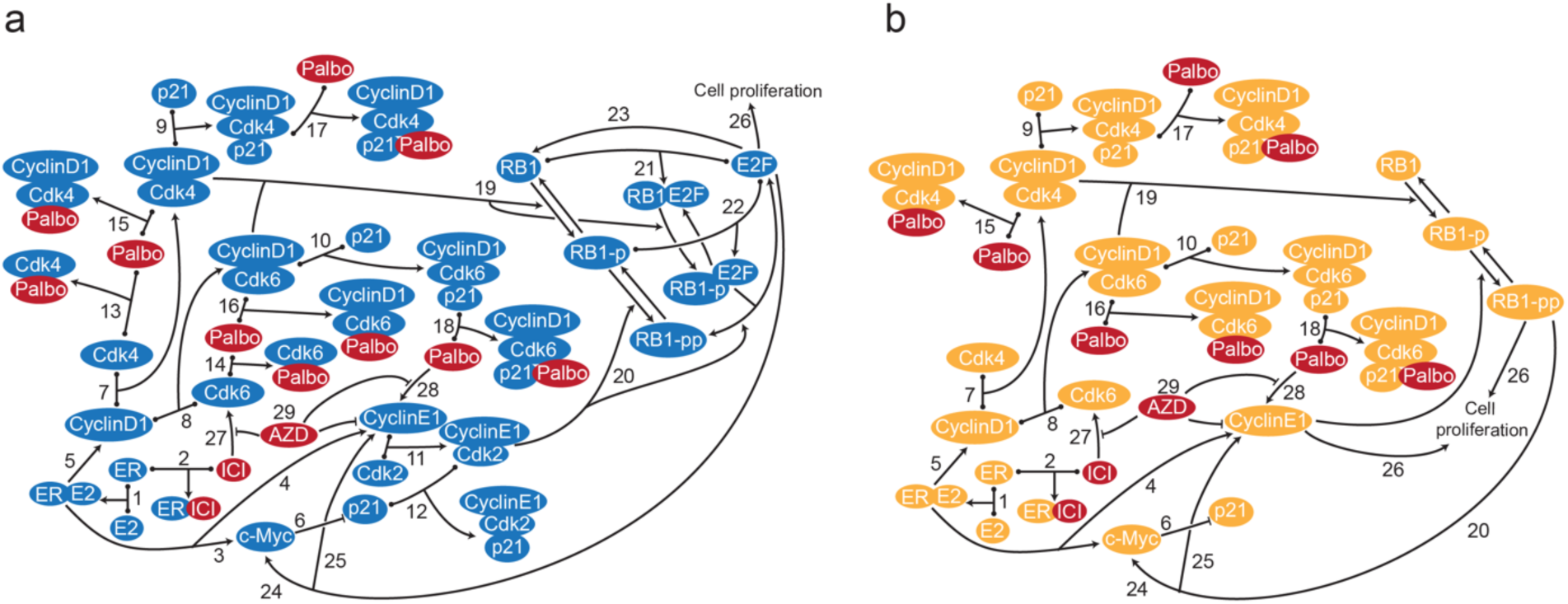
Signaling diagram of the biological mechanism and model structure. **a** Interactions of the biological mechanism. Reversible binding interactions are represented by dots on the components and an arrow to the complex. Arrows pointing from one protein to another protein represent phosphorylation, dephosphorylation, enhancement (arrow) or inhibition (blunt head) of the protein. Line pointing to another line represents enhancement (arrow) or inhibition (blunt head) of the interactions. Applied drugs are colored in red. The biological mechanism consists of the following numbered processes: 1. E2 binds to ER; 2. ICI binds to ER; 3. E2:ER increases transcription of c-Myc; 4. E2:ER increases transcription of CyclinE1; 5. E2:ER increases transcription of CyclinD1; 6. c-Myc inhibits transcription of p21; 7. CyclinD1 binds to Cdk4; 8. CylinD1 binds to Cdk6; 9. p21 binds to CyclinD1:Cdk4; 10. p21 binds to CyclinD1:Cdk6; 11. CyclinE1 binds to Cdk2; 12. p21 binds to CyclinE1:Cdk2; 13. Palbociclib binds to Cdk4; 14. Palbociclib binds to Cdk6; 15. Palbociclib binds to CyclinD1:Cdk4; 16. Palbociclib binds to CyclinD1:Cdk6; 17. Palbociclib binds to CyclinD1:Cdk4:p21; 18. Palbociclib binds to CyclinD1:Cdk6:p21; 19. CyclinD1:Cdk4/6 phosphorylates RB1 to RB1-p; 20. CyclinE1:Cdk2 phosphorylates RB1-p to RB1-pp; 21. RB1 binds to E2F; 22. RB1-p binds to E2F; 23. E2F up-regulates RB1; 24. E2F up-regulates c-Myc; 25. E2F up-regulates CyclinE1. 26. E2F drives cell proliferation; 27. ICI increases Cdk6; 28. Palbociclib increases CyclinE1; 29. AZD decreases CyclinE1, the increased CyclinE1 effect caused by palbociclib and the increased Cdk6 effect caused by ICI. **b** Structure of the mathematical model, a simplified version of the biological mechanism in **a**. Palbo: palbociclib, AZD: AZD1775.

Figs. 5a-d compare the model simulations of 12-month proliferation to experimental results for monotherapy and alternating treatments. The model effectively matches the experimental proliferation results for palbociclib monotherapy, ICI monotherapy, palbociclib and ICI alternating treatment, and palbociclib and ICI plus AZD1775 alternating treatment. By incorporating the resistance mechanisms into the model, we captured the gradually increasing growth observed during the treatments not involving AZD. Through the decreased levels of CyclinE1 and Cdk6 in response to AZD1775, the model can capture the reduced cell proliferation outcomes observed when alternating palbociclib and ICI plus AZD1775 over a 12-month period. The model also captures the 1-month cell proliferation results for AZD1775 at 250nM alone and in combination with ICI at 750nM, Figs. 5e and f. At the same time, the model can reproduce the short-term experimental proliferation results for untreated (E2 control), ICI 500nM treatment, palbociclib 500nM treatment and palbociclib 1μM treatment (Supplementary Fig. 1). The model also captures the significant synergism between ICI and palbociclib (Supplementary Fig. 2), where adding small amounts of palbociclib can dramatically reduce cell proliferation^53^. These results may explain why palbociclib in combination with endocrine therapies achieved substantial improvement in survival outcomes in clinical trials and became the first-line choice of treatment for advanced ER+ breast cancer^4^.

**Fig. 5.**
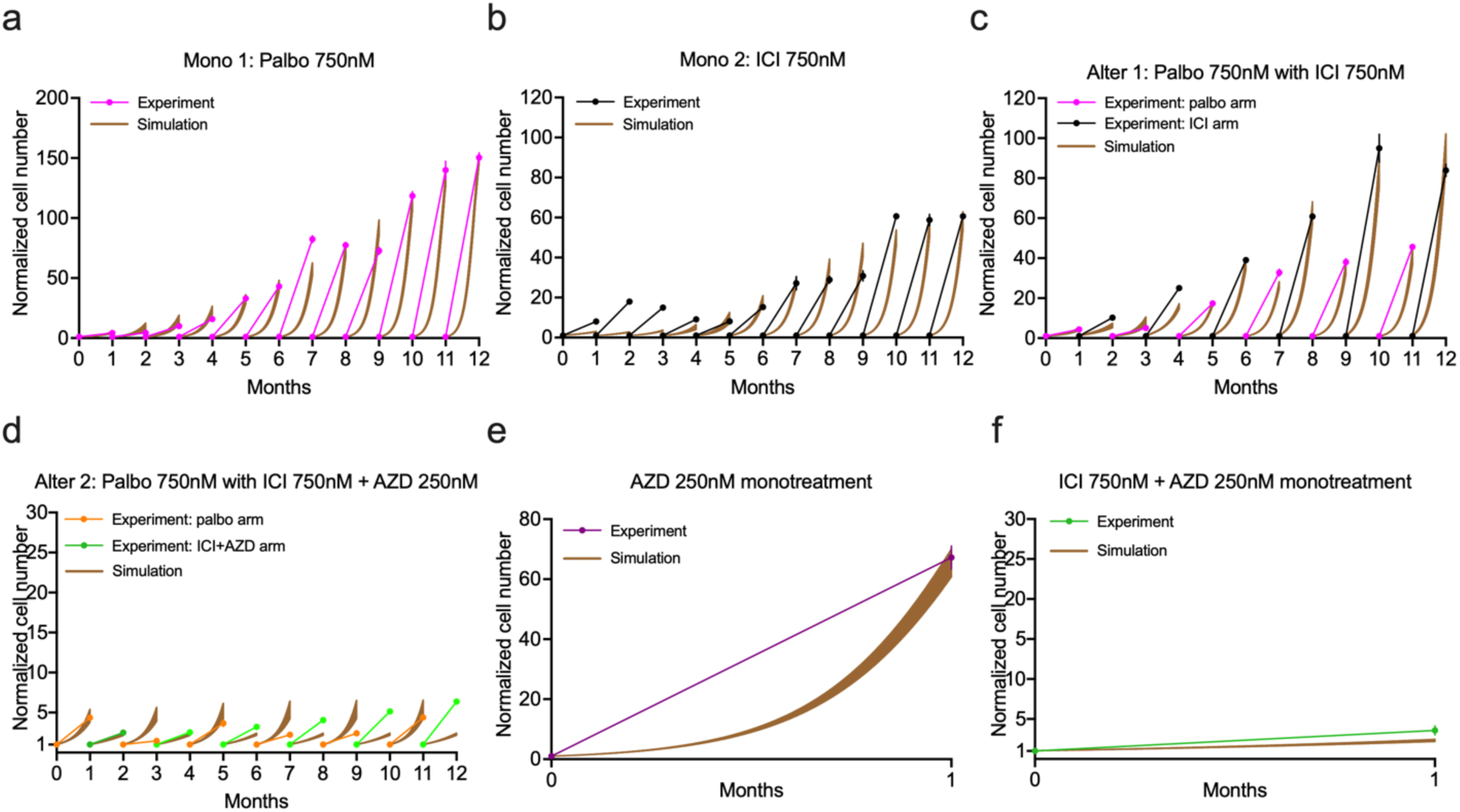
Mathematical model simulation compared to proliferation data for various treatments. **a** Palbociclib monotherapy. The experimental data are shown in magenta and the simulation results are shown in brown (the shaded regions encompass the entire range of simulations within the cohort). **b** ICI monotherapy. The experimental data are shown in black and the simulation results are shown in brown. **c** Palbociclib alternating with ICI treatment. The experimental data are shown in magenta (palbociclib arm) and black (ICI arm). **d** Palbociclib alternating with ICI plus AZD1775 treatment. The experimental data are shown in magenta (palbociclib arm) and green (ICI plus AZD1775 arm). **e** AZD1775 treatment for 1 month. The experimental data are shown in purple. **f** ICI plus AZD1775 treatment for 1 month. The experimental data are shown in green. Palbo: palbociclib, AZD: AZD1775.

### The proposed mathematical model captures the experimental protein data

In the mathematical model, we utilized upregulation of CyclinE1 as the resistance mechanism for palbociclib and upregulation of Cdk6 as the resistance mechanism for ICI. Figs. 6a and b plot the simulation results for total Cdk6 and CyclinE1 levels under the ICI and palbociclib monotherapies, respectively. We see that the total levels of Cdk6 and CyclinE1 slowly increase following ICI and palbociclib monotherapies, respectively. The gradual upregulation of Cdk6 and CyclinE1 mitigates the inhibitory effects of the drugs on cancer cells, allowing them to gradually resume proliferation. In Figs. 6c and d, we see that Cdk6 and CyclinE1 levels are relatively controlled under the two alternating treatments. The reason is that the alternating treatments involve the sequential application of palbociclib and ICI over time, rather than continuously stimulating one particular resistance mechanism. Additionally, by incorporating the resistance suppression effect of AZD1775 on palbociclib and ICI treatment in the model, Cdk6 and CyclinE1 levels are lower in the palbociclib alternating with ICI plus AZD1775 treatment compared to alternating with ICI alone. In Figs. 6e and f, we compared the simulation of Cdk6 and CyclinE1 levels to experimental data under various treatments at the experimental timepoints. The model simulation results align closely with the protein levels measured in the experiments.

**Fig. 6.**
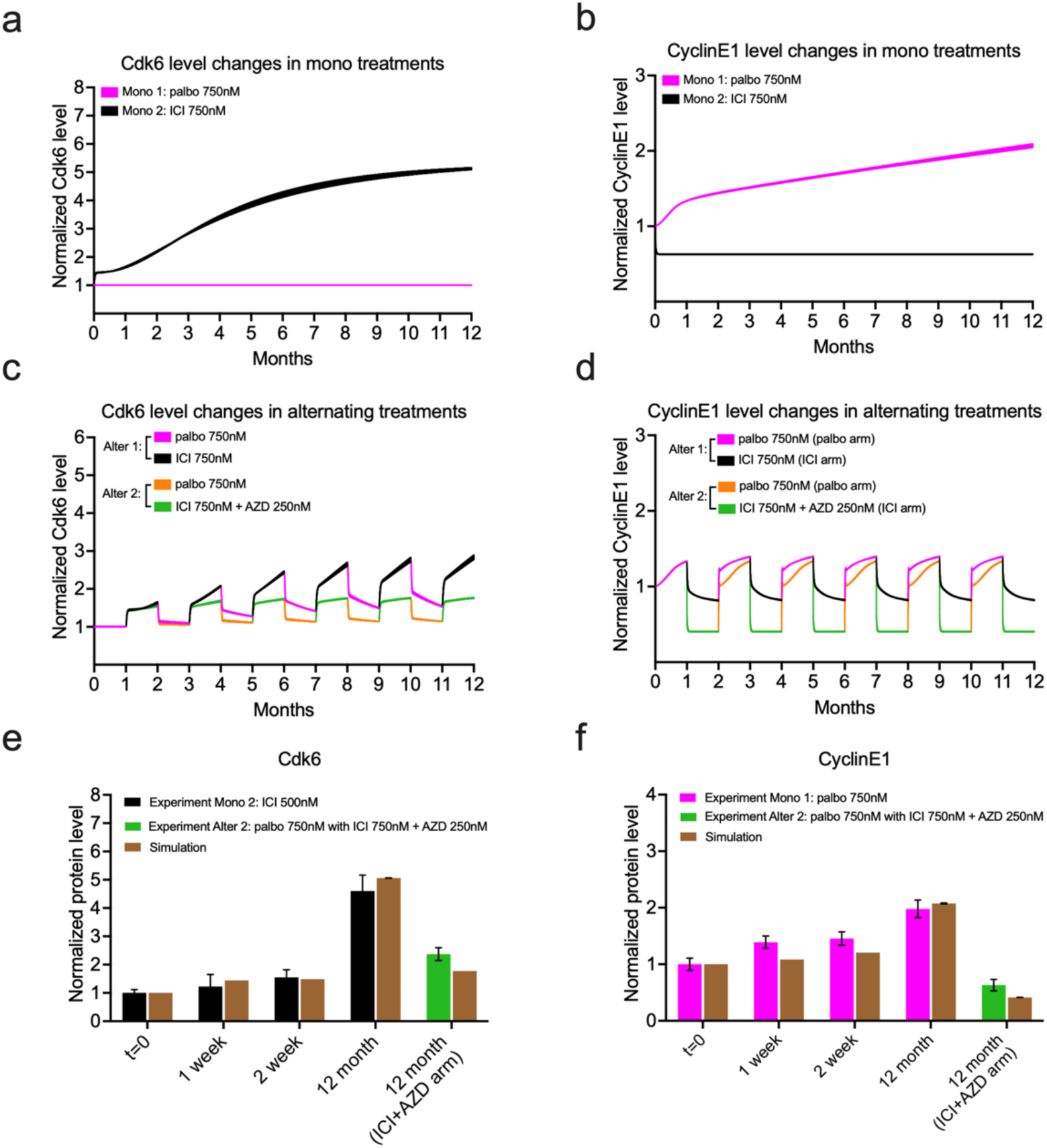
Mathematical model simulation of protein level changes and comparison to western blot data for various treatments. **a** Model simulation of normalized Cdk6 level changes for palbociclib (magenta) and ICI (black) monotherapy over 12 months. The shaded regions encompass the entire range of simulations for the cohort. **b** Model simulation of normalized CyclinE1 level changes for palbociclib (magenta) and ICI (black) monotherapy over 12 months. **c** Model simulation of normalized Cdk6 level changes for palbociclib (magenta) alternating with ICI (black), and palbociclib (orange) alternating with ICI plus AZD1775 (green) over 12 months. **d** Model simulation of normalized CyclinE1 level changes for palbociclib (magenta) alternating with ICI (black), and palbociclib (orange) alternating with ICI plus AZD1775 (green) over 12 months. **e** Bar plot of model simulation of Cdk6 level compared to experimental data for ICI (black) monotherapy, and palbociclib alternating with ICI plus AZD1775 (green) at different timepoints. The simulation results shown in brown are the average results from all cohort simulations. **f** Bar plot of model simulation of CylinE1 level compared to experimental data for palbociclib (magenta) monotherapy, and palbociclib alternating with ICI plus AZD1775 (green) at different timepoints. Palbo: palbociclib, AZD: AZD1775.

### Optimal treatment design using the model

One of the key benefits of a mathematical model is that it enables us to explore a wide range of treatment options to find the best possibilities. “Best” might be defined as maximizing tumor regression or as minimizing the toxicity and side effects associated with treatment. In our scenario, where the side effects of AZD1775 are a significant concern, we used the mathematical model to identify alternating treatment regimens that minimize the total duration of ICI plus AZD1775 periods while still maintaining its efficacy. Figs. 7a-c show a proposed palbociclib and ICI plus AZD1775 alternating treatment which has the minimum frequency of ICI plus AZD1775 treatment intervals while ensuring that the maximum proliferation does not exceed 20-fold per month within 24 months. As shown in Fig. 7a, the proposed alternating treatment consists of a total of 6 arms of ICI plus AZD1775 treatment. Before 16 months, one month of ICI plus AZD1775 treatment is necessary every three months of palbociclib treatment to prevent the development of palbociclib resistance. As time progresses, the gradual increase in CyclinE1 levels during palbociclib treatment necessitates ICI plus AZD1775 treatment every two months of palbociclib therapy. In Figs. 7b and c, we see that total CyclinE1 and Cdk6 levels are effectively controlled, resulting in the suppression of cell proliferation throughout the entire alternating treatment period.

**Fig. 7.**
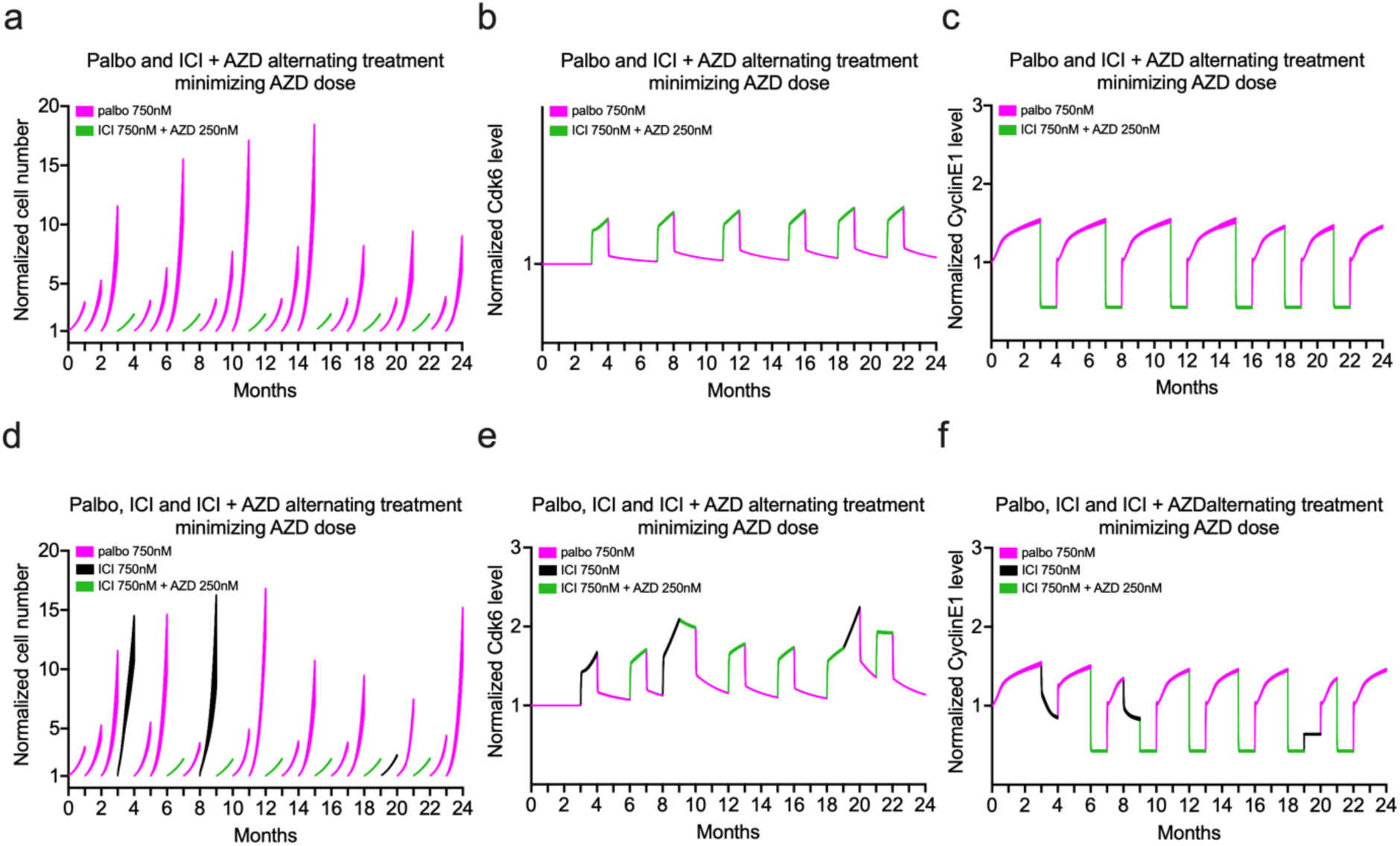
Minimize the number of ICI plus AZD1775 treatment intervals within an alternating regimen while ensuring that the maximum proliferation does not exceed 20-fold per month over 24 months. **a-c** Proposed Palbociclib (magenta) alternating with ICI plus AZD1775 (green) protocol to reduce the frequency of ICI plus AZD1775. The shaded regions encompass the entire range of simulations within the cohort. **a** Normalized cell number. **b** Normalized total Cdk6 level. **c** Normalized total CyclinE1 level. **d**-**f** Proposed Palbociclib (magenta), ICI (black) and ICI plus AZD1775 (green) alternating treatment. **d** Normalized cell number. **e** Normalized total Cdk6. **f** Normalized total CyclinE1. Palbo: palbociclib, AZD: AZD1775.

Figs. 7d and e present another proposed alternating treatment with three options: palbociclib alone, ICI alone or ICI plus AZD1775. The treatment is also designed to minimize the frequency of ICI plus AZD1775 treatment intervals while ensuring that the maximum number of cell proliferations does not exceed 20-fold per month within a 24-month period. As shown in Figs. 7d and e, the proposed alternating treatment comprises a total of 6 arms of ICI plus AZD1775 treatment. The total CyclinE1 and Cdk6 level are effectively regulated, leading to the suppression of cell proliferation throughout the entire alternating treatment period without developing significant resistance to palbociclib or ICI.

## Discussion

Targeted therapy can become ineffective over time as cancer cells adapt to drugs through various mechanisms, evading the effects of therapy and continuing to grow and spread. Combination and alternating treatment regimens are strategies commonly used to fight resistance. By constantly changing the selective pressure on cancer cells, alternating treatments make it more difficult for resistance mechanisms to fully develop. Combination treatments use two or more drugs with complementary mechanisms to increase treatment efficacy. Cdk4/6 inhibitors in combination with endocrine therapy have emerged as the standard of care for hormone receptor-positive (HR+) metastatic and advanced breast cancer. Ongoing research focusing on preventing the development of resistance to these drugs remains crucial for further improving outcomes for these patients^8,9^. Our work addressed this question by investigating an alternating regimen that involves sequentially applying palbociclib and the combination of ICI and AZD1775. We demonstrated that alternating palbociclib with the combination of ICI plus AZD1775 prevents the development of drug resistance to both palbociclib and ICI for 12 months. Subsequently, we extended our previous mathematical model^52–54^ to recapitulate the new experimental cell proliferation results. We utilized the observed upregulation of Cdk6 and CyclinE1 in western blot data to model the progressive development of palbociclib and ICI resistance, respectively. We incorporated the effects of AZD1775 into the model and calibrated using the protein data, enabling the model to reproduce the effect of alternating treatment delaying the resistance to palbociclib and ICI. Finally, to address the prevalent concern of AZD1775 side effects, we demonstrate that the mathematical model can be utilized to propose a treatment regimen that minimizes the exposure to the drug while effectively controlling the proliferation. One of the most significant advantages of building a mathematical model is to enable exploration of better treatment regimens tailored to different constraints by considering various drug selections, combinations and dosing strategies.

In our experimental results, merely alternating palbociclib with ICI somewhat delayed the development of resistance and allowed the cells to maintain some sensitivity to palbociclib after 12 months of treatment. But the results were not satisfactory in terms of decreasing proliferation over the long term, likely because we continually targeted the G1/S transition. But drug resistance often comes with an associated fitness cost, so resistance to one drug may be associated with an acquired vulnerability to another drug, a phenomenon referred to as “collateral sensitivity”^16^. Alternating treatment of the drug-sensitive and drug-resistant populations can keep the tumor under control over longer periods of time. An example of this is that sequential, but not simultaneous, treatment of triple-negative breast cancer cells with EGFR inhibitors and DNA-damaging drugs leads to efficient cell killing^58^. We believe that MCF7 cells heavily rely on their ability to manage the higher levels of replication stress caused by the increased kinase activities creating resistance. This elevated reliance exacerbates their vulnerability to WEE1 inhibition and accounts for the success of adding AZD1775 to the alternating regimen. Identification of collateral sensitivities of drug-resistant cancer cells may pave the way for innovative treatment regimens that delay or prevent the emergence of drug resistance.

Finally, we acknowledge the limitations of our current work. The absence of an AZD1775 resistance mechanism in the model could impede the exploration of optimized treatments over extended periods^59^. Additionally, the reliance on results solely from the MCF7 cell line for the alternating treatment likely does not fully represent the diversity of ER+ breast cancer cells. Further testing on other ER+ breast cancer cell lines, such as the T47D cell line, will be necessary to generalize our findings. But we remain optimistic that uncovering the vulnerabilities of resistant cells can pave the way for alternating treatments that target both drug-sensitive and drug-resistant populations in a periodic fashion, ultimately enabling the control of tumors over prolonged periods of time and delivering long-lasting benefits to patients.

## Authors’ Contributions

W. H.: Resources, data curation, software, formal analysis, validation, investigation, visualization, methodology, writing-original draft, writing-review and editing. D.M.D.: Data curation, investigation. P. K.: Conceptualization, supervision, methodology, writing-review and editing. A.N. S.-H.: Conceptualization, resources, supervision, funding acquisition, investigation, methodology, writing-review and editing. W.T. B.: Conceptualization, resources, supervision, funding acquisition, investigation, methodology, writing-review and editing.

## Materials and Methods

### Cell culture and drugs

MCF7 cells (RRID:CVCL_0031) were obtained from Tissue Culture Shared Resources at Lombardi Comprehensive Cancer Center, Georgetown University, Washington, DC. MCF7 cells were grown in phenol red-free improved minimal essential medium (Life Technologies, Grand Island, NY; A10488-01) with 10% charcoal-stripped calf serum (CCS) and supplemented with 10nM 17b-estradiol (E2). ICI and palbociclib were obtained from Tocris Bioscience (Ellisville, MO). AZD1775 were obtained from Cayman Chemical (Ann Arbor, MI). MCF7 cells were authenticated by DNA fingerprinting and tested regularly for Mycoplasma infection. All other chemicals were purchased from Sigma-Aldrich (St. Louis, MO). We used MCF7 cells in our experiments that was derived from a female patient. Given that ER+ breast cancer is a malignancy that occurs overwhelmingly in females, this is scientifically justified.

### Cell proliferation assay

MCF7 cells were seeded at a density of 4-5 × 10^4^ cells/well in 60mm plates and treated with indicated drugs at 24-hour post plating. To measure cell number at number at specific time-points, cells were trypsinized, resuspended in phosphate-buffered saline (PBS) and counted using a Z1 Single Coulter Counter (Beckman Coulter, Miami, FL).

### Western blot analysis

For Western blot analysis, cells were lysed for 30 minutes on ice with lysis buffer (50 mM Tris-HCI, pH 7.5, containing 150nM NaCl, 1mM EDTA, 0.5% sodium deoxycholate, 1% IGEPAL CA-630, 0.1% sodium dodecylsulfate (SDS), 1mM Na3VO4, 44 μg ml–1 phenylmethylsulfonyl fluoride) supplemented with Complete Mini protease inhibitor mixture tablets (Roche Applied Science). Total protein was quantified using the bicinchoninic acid assay (Pierce). Whole-cell lysate (20 mg) was resolved by SDS–polyacrylaminde gel electrophoresis.

### Parameter calibration of the mathematical model

The biological interactions we considered are based on known mechanisms from the literature. Since we ignored many other (known and unknown) interactions that affect these transitions, it is unlikely that our model can exactly fit the data. So, in addition to the best-fit parameter set, we created a cohort of 99 additional parameter sets (total 100 parameter sets) that fit the data only slightly less well than the optimal set (increased squared deviation between experiment and simulation less than about 20% of the optimal). In the figures that follow, we plot the range of results from simulating the entire cohort. The best-fit parameter set was calibrated using the *patternsearch* function in MATLAB (RRID:SCR_001622, R2023a) to reduce the least-squares difference between the model simulation and the experimental results^52,53^. The parameter cohort was generated using the *ga* function in MATLAB. The mathematical model contains 26 ordinary differential equations (ODEs) and 93 parameters (75 calibrated and 18 fixed, shown in the Supplementary file), which is implemented in MATLAB. The generation, degradation, phosphorylation, dephosphorylation, binding and unbinding reactions are modeled by mass action laws and hill functions. Drug treatment effects are modeled by competitive binding to their targets. The ODEs are solved numerically by the ode23tb function in MATLAB. Code for the model is available at (https://github.com/weihevt/WEE1Cdk46).

### Mathematical model structure

The structure of our ODE model is based on the signaling pathway of the G1-S transition since palbociclib and ICI primarily affect progression through the G1 phase of the cell cycle. It is impractical to include all interactions related to drug treatments in the biological mechanism, so we retained only those necessary to capture the main effects of the treatments (see Results). The justifications for the interactions we included are listed in the Supplementary file. This biological model structure was further simplified to ease mathematical modeling. The major modification is not including E2F in the model and instead using RB1-pp and CyclinE1 to represent E2F transcriptional activity and govern the proliferation rate^52–54^.

## Data Availability

Data used to generate the figures is available at https://github.com/weihevt/WEE1Cdk46.

## Code Availability

Code is included in the electronic supplementary material and is also available at https://github.com/weihevt/WEE1Cdk46.

## Acknowledgments

This work was partly supported by Public Health Service grant R01-CA201092 to W.T.B. and A.N.S.-H. Technical services were provided by Shared Resources at Georgetown University Medical Center, including the Tissue Culture Core Shared Resources and the Genomics and Epigenomics Shared Resources, that were funded through Public Health Service award 1P30-CA-51008 (Lombardi Comprehensive Cancer Center Support Grant). We also thank the Georgetown Breast Cancer Advocates (GBCA) for a patient’s perspective for this study.

