## Supplemental for "WEE1 inhibition delays resistance to CDK4/6 inhibitor and antiestrogen treatment in estrogen receptor-positive breast cancer"

### Supplementary Materials and Methods:

#### Biological signaling diagram

The literature references for the numbered interactions of the signaling pathway and drugs, shown in Fig. 3a, are as follows:

1. E2 binds to ER[1]; 2. ICI binds to ER[2]; 3. E2:ER increases transcription of c-Myc[3]; 4. E2:ER increases transcription of cyclinE1[4]; 5. E2:ER increases transcription of cyclinD1[3]; 6. c-Myc inhibits transcription of p21[5]; 7. CyclinD1 binds to Cdk4[6]; 8. CylinD1 binds to Cdk6[6]; 9. p21 binds to cyclinD1:Cdk4[7]; 10. p21 binds to cyclinD1:Cdk6[7]; 11. CyclinE1 binds to Cdk2[3]; 12. p21 binds to cyclinE1:Cdk2[8]; 13. Palbociclib binds to Cdk4[9]; 14. Palbociclib binds Cdk6[9]; 15. Palbociclib binds to cyclinD1:Cdk4[9]; 16. Palbociclib binds to cyclinD1:Cdk6[9]; 17. Palbociclib binds to cyclinD1:Cdk4:p21[10]; 18. Palbociclib binds to cyclinD1:Cdk6:p21[10]; 19. CyclinD1:Cdk4/6 phosphorylates RB1 to RB1-p[3]; 20. CyclinE1:Cdk2 phosphorylates RB1-p to RB1-pp[3]; 21. RB1 binds to E2F[11]; 22. RB1-p binds to E2F[11]; 23. E2F up-regulates RB1[12]; 24. E2F up-regulates c-Myc[13]; 25. E2F up-regulates cyclinE1[14]. 26. E2F drives cell proliferation[15]; 27. ICI increases Cdk6[16–18]; 28. Palbociclib increases cyclinE1[19–24]; 29. AZD decreases cyclinE1, the cyclinE1 increase effect caused by palbociclib and the Cdk6 increase effect caused by ICI[25–28].

#### Model Variables

| Variable name | Description | Half-life |
|---|---|---|
| (1) $E2_{media}$ | E2 concentration in the media | - |
| (2) $E2_{cell}$ | E2 concentration in the cell | - |
| (3) *ER* | Estrogen receptor α | ~4-5h[29] |
| (4) *E2ER* | Estrogen bound estrogen receptor α | ~3-4h[29] |
| (5) *ICIER* | ICI 182,780 bound estrogen receptor | < 3-4h[29] |
| (6) *rescyclinE1palbo* | Variable induced by palbociclib increasing cyclinE1 | - |
| (7) rescdk6ICI | Variable induced by ICI increasing cdk6 | - |
| (8) *cyclinD1* | cyclinD1 | ~0.4h[30] |
| (9) *cdk4* | Cdk4 | ~5h[31] |
| (10) *cdk6* | Cdk6 | ~5h[31] |
| (11) *cyclinD1cdk4* | CyclinD1 bound Cdk4 | - |
| (12) *cyclinD1cdk6* | CyclinD1 bound Cdk6 | - |
| (13) *cyclinD1cdk4p21* | p21 bound cyclinD1:Cdk4 | - |
| (14) *cyclinD1cdk6p21* | p21 bound cyclinD1:Cdk6 | - |
| (15) *cyclinD1cdk4palbo* | Palbociclib bound cyclinD1:Cdk4 | - |
| (16) *cyclinD1cdk6palbo* | Palbociclib bound cyclinD1:Cdk6 | - |
| (17) *cyclinD1cdk4p21palbo* | Palbociclib bound cyclinD1:Cdk4:p21 | - |
| (18) *cyclinD1cdk6p21palbo* | Palbociclib bound cyclinD1:Cdk6:p21 | - |
| (19) *cMyc* | c-Myc | ~0.3h[32] |
| (20) *p21* | p21 | ~0.3-1h[33] |

| | | |
|---|---|---|
| (21) *cyclinE1* | cyclinE1 | ~0.5h[34] |
| (22) *Rb* | Retinoblastoma protein | ~2-3h[35] |
| (23) *pRb* | Hypophosphorylated RB1 (RB1-p) | ~2-3h[35](35) |
| (24) *ppRb* | Hyperphosphorylated RB1 (RB1-pp) | > 4h[35] |
| (25) *Nalive* | Alive cell number | - |
| (26) *Ndead* | Dead cell number | - |

#### Model Parameter Descriptions, Values and Declaration of Fixed or Calibrated

| Parameter name | Description | Value | Fixed/ Calibrated |
|---|---|---|---|
| (1) $k_{diff}$ | Diffusion rate of E2 | 3.37/h | Calibrated |
| (2) $Vol_{1cell}$ | Volume of MCF7 cell | $8\times10^{-5}$mL | Fixed |
| (3) $Vol_{media}$ | Volume of media | 10mL | Fixed |
| (4) $k_{ER}$ | Translation rate of *ER* | 316.28nM/h | Calibrated |
| (5) $kd_{ER}$ | Degradation rate of *ER* | 0.1/h | Fixed |
| (6) $kd_{E2ER}$ | Degradation rate of *E2ER* | 0.3/h | Fixed |
| (7) $kd_{ICIER}$ | Degradation rate of *ICIER* | 4.81/h | Calibrated |
| (8) $kb_{E2ER}$ | Binding rate between *ER* and $E2_{cell}$ | 143.73/(h×nM) | Calibrated |
| (9) $kub_{E2ER}$ | Unbinding rate between *ER* and $E2_{cell}$ | 1.0/h | Fixed |
| (10) $kb_{ICIER}$ | Binding rate between *ICI* and *ER* | 0.3/(h×nM) | Calibrated |
| (11) $kub_{ICIER}$ | Unbinding rate between *ICI* and *ER* | 1.0/h | Fixed |
| (12) $k_{cyclinD1}$ | Translation rate of *cyclinD1* | 11.33nM/h | Calibrated |
| (13) $kd_{cyclinD1}$ | Degradation rate of *cyclinD1* | 1.4/h | Fixed |
| (14) $kb_{cyclinD1cdk46}$ | Binding rate between *cyclinD1* and *cdk46* | $1.7\times10^{4}$ | Calibrated |
| (15) $kub_{cyclinD1cdk46}$ | Unbinding rate between *cyclinD1* and *cdk46* | 1.0/h | Fixed |
| (16) $k_{cyclinD1E2ER}$ | Increasing rate of *cyclinD1* by *E2ER* | 349.12nM/h | Calibrated |
| (17) $p_{cyclinD1E2ER_1}$ | Parameter 1 of *cyclinD1* increased by *E2ER* | 1592.05nM | Calibrated |
| (18) $p_{cyclinD1E2ER_2}$ | Parameter 2 of *cyclinD1* increased by *E2ER* | 9.03 | Calibrated |
| (19) $k_{rescdk6ICI}$ | Increasing rate of *rescdk6ICI* | $6.98\times10^{-5}$nM/h | Calibrated |
| (20) $p_{rescdk6ICI_1}$ | Parameter 1 of *rescdk6ICI* increased by *ICI* | 1380.8nM | Calibrated |
| (21) $p_{rescdk6ICI_2}$ | Parameter 2 of *rescdk6ICI* increased by *ICI* | 3.25 | Calibrated |
| (22) $kd_{rescdk6ICI}$ | Decreasing rate of *rescdk6ICI* | $1.66\times10^{-5}$/h | Calibrated |
| (23) $p_{kdrescdk6ICI_1}$ | Parameter 1 of *rescdk6ICI* decreasing | 0.027nM | Calibrated |
| (24) $p_{kdrescdk6ICI_2}$ | Parameter 2 of *rescdk6ICI* decreasing | 1.29 | Calibrated |
| (25) $kd_{rescdkICI_{AZD}}$ | Decreasing rate of *rescdk6ICI* by *AZD* | $9.02\times10^{-4}$/h | Calibrated |
| (26) $k_{cdk4}$ | Translation rate of *cdk4* | 18.23nM/h | Calibrated |
| (27) $kd_{cdk46}$ | Degradation rate of *cdk46* | 0.1155/h | Fixed |
| (28) $k_{cdk6}$ | Translation rate of *cdk6* | 0.2nM/h | Calibrated |
| (29) $k_{cdk6rescdk6ICI}$ | Increasing rate of *cdk6* by *rescdk6ICI* | 1.45nM/h | Calibrated |
| (30) $p_{cdk6rescdk6ICI_1}$ | Parameter 1 of *cdk6* increased by *rescdk6ICI* | 0.035nM | Calibrated |
| (31) $p_{cdk6rescdk6ICI_2}$ | Parameter 2 of *cdk6* increased by *rescdk6ICI* | 1.93 | Calibrated |
| (32) $kd_{cyclinD1cdk46}$ | Degradation rate of *cyclinD1cdk46* | 0.19/h | Calibrated |

| | | | |
|---|---|---|---|
| (33) $kb_{cyclinD1cdk46palbo}$ | Binding rate between *cyclinD1cdk46* and *palbo* | 0.0083/(h×nM) | Calibrated |
| (34) $kub_{cyclinD1cdk46palbo}$ | Unbinding rate between *cyclinD1cdk46* and *palbo* | 1.0/h | Fixed |
| (35) $kb_{cyclinD1cdk46p21}$ | Binding rate between *cyclinD1cdk46* and *p21* | 4.13/(h×nM) | Calibrated |
| (36) $kub_{cyclinD1cdk46p21}$ | Unbinding rate between *cyclinD1cdk46* and *p21* | 1.0/h | Fixed |
| (37) $kb_{cyclinD1cdk46p21palbo}$ | Binding rate between *cyclinD1cdk46p21* and *palbo* | 14.51/(h×nM) | Calibrated |
| (38) $kub_{cyclinD1cdk46p21palbo}$ | Unbinding rate between *cyclinD1cdk46p21* and *palbo* | 1.0/h | Fixed |
| (39) $k_{cMyc}$ | Translation rate of *cMyc* | 13.33nM/h | Calibrated |
| (40) $kd_{cMyc}$ | Degradation rate of *cMyc* | 2.31/h | Fixed |
| (41) $k_{cMycE2ER}$ | Increasing rate of *cMyc* by *E2ER* | 417.36nM/h | Calibrated |
| (42) $p_{cMycE2ER_1}$ | Parameter 1 of *cMyc* increased by *E2ER* | 1811.48nM | Calibrated |
| (43) $p_{cMycE2ER_2}$ | Parameter 2 of *cMyc* increased by *E2ER* | 5.59 | Calibrated |
| (44) $k_{cMycppRb}$ | Increasing rate of *cMyc* by *ppRb* | 59.93nM/h | Calibrated |
| (45) $p_{cMycppRb_1}$ | Parameter 1 of *cMyc* increased by *ppRb* | 18.13nM | Calibrated |
| (46) $p_{cMycppRb_2}$ | Parameter 2 of *cMyc* increased by *ppRb* | 3.76 | Calibrated |
| (47) $k_{p21cMyc}$ | Translation rate of *p21* inhibited by *cMyc* | 11.21nM/h | Calibrated |
| (48) $p_{p21cMyc_1}$ | Parameter 1 of *p21* inhibited by *cMyc* | 6.85nM | Calibrated |
| (49) $p_{p21cMyc_2}$ | Parameter 2 of *p21* inhibited by *cMyc* | 4.24 | Calibrated |
| (50) $kd_{p21}$ | Degradation rate of *p21* | 1.39/h | Fixed |
| (51) $k_{rescyclinE1palbo}$ | Increasing rate of *rescyclinE1palbo* | $4.5\times10^{-4}$nM/h | Calibrated |
| (52) $p_{rescyclinE1palbo_1}$ | Parameter 1 of *rescyclinE1palbo* increased by *palbo* | 197.66nM | Calibrated |
| (53) $p_{rescyclinE1palbo_2}$ | Parameter 2 of *rescyclinE1palbo* increased by *palbo* | 1.19 | Calibrated |
| (54) $kd_{rescyclinE1palbo}$ | Decreasing rate of *rescyclinE1palbo* | $3.34\times10^{-4}$/h | Calibrated |
| (55) $p_{kdrescyclinE1palbo_1}$ | Parameter 1 of *rescyclinE1palbo* decreasing | 0.14nM | Calibrated |
| (56) $p_{kdrescyclinE1palbo_2}$ | Parameter 2 of *rescyclinE1palbo* decreasing | 10 | Calibrated |
| (57) $kd_{rescyclinE1palbo_{AZD}}$ | Decreasing rate of *rescyclinE1palbo* by *AZD* | 7.64/h | Calibrated |
| (58) $k_{cyclinE1}$ | Translation rate of *cyclinE1* | 0.39nM/h | Calibrated |
| (59) $kd_{cyclinE1}$ | Degradation rate of *cyclinE1* | 1.39/h | Fixed |
| (60) $kd_{cyclinE1_{AZD}}$ | Decreasing rate of *cyclinE1* by *AZD* | 0.66/h | Fixed |
| (61) $k_{cyclinE1E2ER}$ | Increasing rate of *cyclinE1* by *E2ER* | 1.39nM/h | Calibrated |
| (62) $p_{cyclinE1E2ER_1}$ | Parameter 1 of *cyclinE1* increased by *E2ER* | 892.32nM | Calibrated |
| (63) $p_{cyclinE1E2ER_2}$ | Parameter 2 of *cyclinE1* increased by *E2ER* | 2.92 | Calibrated |
| (64) $k_{cyclinE1rescyclinE1palbo}$ | Increasing rate of *cyclinE1* by *rescyclinE1palbo* | 5.76nM/h | Calibrated |
| (65) $p_{cyclinE1rescyclinE1palbo_1}$ | Parameter 1 of *cyclinE1* increase by *rescyclinE1palbo* | 1.03nM | Calibrated |
| (66) $p_{cyclinE1rescyclinE1palbo_2}$ | Parameter 2 of *cyclinE1* increase by *rescyclinE1palbo* | 1.36 | Calibrated |
| (67) $k_{Rb}$ | Translation rate of *Rb* | 10.44nM/h | Calibrated |
| (68) $kd_{Rb}$ | Degradation rate of *Rb* | 0.35/h | Fixed |
| (69) $k_{RbppRb}$ | Increasing rate of *Rb* by *ppRb* | 7.13nM/h | Calibrated |
| (70) $p_{RbppRb_1}$ | Parameter 1 of *Rb* increased by *ppRb* | 1.55nM | Calibrated |
| (71) $p_{RbppRb_2}$ | Parameter 2 of *Rb* increased by *ppRb* | 8.18 | Calibrated |
| (72) $k_{RbcyclinD1cdk4}$ | Phosphorylation rate of *Rb* by *cyclinD1cdk4* | 0.25/h | Calibrated |
| (73) $p_{cyclinD1cdk4_1}$ | Parameter 1 of *cyclinD1cdk4* kinase activity | 826.77nM | Calibrated |
| (74) $p_{cyclinD1cdk4_2}$ | Parameter 2 of *cyclinD1cdk4* kinase activity | 0.17 | Calibrated |
| (75) $k_{RbcyclinD1cdk6}$ | Phosphorylation rate of *Rb* by *cyclinD1cdk6* | 1.28/h | Calibrated |

| | | | |
|---|---|---|---|
| (76) $p_{cyclinD1cdk6_1}$ | Parameter 1 of *cyclinD1cdk6* kinase activity | 1.91nM | Calibrated |
| (77) $p_{cyclinD1cdk6_2}$ | Parameter 2 of *cyclinD1cdk6* kinase activity | 1.27 | Calibrated |
| (78) $k_{pRbcyclinE1}$ | Phosphorylation rate of *pRb* by *cyclinE1* | 84.2/h | Calibrated |
| (79) $p_{cyclinE1_1}$ | Parameter 1 of *cyclinE1* kinase activity | 1.22nM | Calibrated |
| (80) $p_{cyclinE1_2}$ | Parameter 2 of *cyclinE1* kinase activity | 1.97 | Calibrated |
| (81) $k_{pRbdepho}$ | Dephosphorylation rate of *pRb* | 6.5nM/h | Calibrated |
| (82) $kd_{ppRb}$ | Degradation rate of *ppRb* | 0.05/h | Fixed |
| (83) $k_{ppRbdepho}$ | Dephosphorylation rate of *ppRb* | 21.89nM/h | Calibrated |
| (84) $k_{pro}$ | Basal proliferation rate | 0.0016 | Calibrated |
| (85) $k_{proppRb}$ | Proliferation rate increased by *ppRb* | 0.04 | Calibrated |
| (86) $p_{proppRb_1}$ | Parameter 1 of proliferation rate increased by *ppRb* | 4.27nM | Calibrated |
| (87) $p_{proppRb_2}$ | Parameter 2 of proliferation rate increased by *ppRb* | 3.3 | Calibrated |
| (88) $k_{procyclinE1}$ | Proliferation rate increased by *cyclinE1* | 8.16 | Calibrated |
| (89) $p_{procyclinE1_1}$ | Parameter 1 of proliferation rate increased by *cyclinE1* | 5.27nM | Calibrated |
| (90) $p_{procyclinE1_2}$ | Parameter 2 of proliferation rate increased by *cyclinE1* | 7.41 | Calibrated |
| (91) $k_{carrying}$ | Carrying capacity | 160.31 | Calibrated |
| (92) $k_{death}$ | Death rate | $2.65\times10^{-4}$/hour | Calibrated |
| (93) $k_{lysis}$ | Lysis rate of dead cell | $2.12\times10^{-4}$/hour | Calibrated |
| (94) *E2* | $E2_{media}$ in control condition | 10nM | Fixed |
| (95) *ICI* | Concentration of ICI 182,780 | Varies | Fixed |
| (96) *palbo* | Concentration of palbociclib | Varies | Fixed |
| (97) *AZD* | Concentration of AZD1775 | 250nM | Fixed |

**Model Equations**

$$N = N_{alive} + N_{dead} \tag{1}$$

(1) Total number of cells equals number of alive cells plus number of dead cells

$$\frac{dN_{alive}}{dt} = (k_{pro} + k_{procyclinE1} \times \frac{cyclinE1^{p_{procyclinE1_2}}}{p_{procyclinE1_1}^{p_{procyclinE1_2}} + cyclinE1^{p_{procyclinE1_2}}} + k_{proppRb} \times \frac{ppRb^{p_{proppRb_2}}}{p_{proppRb_1}^{p_{proppRb_2}} + ppRb^{p_{proppRb_2}}}) \times N_{alive} \times (1 - \frac{N_{alive}}{k_{carrying}}) \tag{2}$$

$$-k_{death} \times N_{alive} \tag{3}$$

(2) Basal proliferation, increased proliferation by $cyclinE1$ and $ppRb$

(3) Basal death

$$\frac{dN_{dead}}{dt} = k_{death} \times N_{alive} \tag{4}$$

$$-k_{lysis} \times N_{dead} \tag{5}$$

(4) Basal death

(5) Lysis of dead cells

$$\frac{dE2_{media}}{dt} = \frac{k_{diff} \times N \times Vol_{1cell}}{Vol_{media}} \times (E2_{cell} - E2_{media}) \quad (6)$$

(6) E2 concentration changes in media

$$\frac{dE2_{cell}}{dt} = -k_{diff} \times (E2_{cell} - E2_{media}) - \frac{\left(\frac{dN_{alive}}{dt} + \frac{dN_{dead}}{dt}\right)}{N} \times E2_{cell} \quad (7)$$

$$-kb_{E2ER} \times E2_{cell} \times ER + kub_{E2ER} \times E2ER \quad (8)$$

$$-kb_{NSB} \times E2_{cell} + kub_{NSB} \times E2NSB \quad (9)$$

$$+\, kd_{E2ER} \times E2ER \quad (10)$$

(7) E2 concentration changes in cell

(8) Binding and unbinding between $ER$ and $E2_{cell}$

(9) Binding and unbinding between non-specific binding and $E2_{cell}$ in the cell

(10) Degradation of $E2ER$

$$\frac{dER}{dt} = k_{ER} - kd_{ER} \times ER \quad (11)$$

$$-kb_{E2ER} \times E2_{cell} \times ER + kub_{E2ER} \times E2ER \quad (12)$$

$$-kb_{ICIER} \times ICI \times ER + kub_{ICIER} \times ICIER \quad (13)$$

(11) Translation and degradation of $ER$

(12) Binding and unbinding between $ER$ and $E2_{cell}$

(13) Binding and unbinding between $ER$ and $ICI$

$$\frac{dE2ER}{dt} = -kd_{E2ER} \times E2ER \quad (14)$$

$$+kb_{E2ER} \times E2_{cell} \times ER - kub_{E2ER} \times E2ER \quad (15)$$

(14) Degradation of $E2ER$

(15) Binding and unbinding between $ER$ and $E2_{cell}$

$$\frac{dICIER}{dt} = kb_{ICIER} \times ICI \times ER - kub_{ICIER} \times ICIER \quad (16)$$

$$-kd_{ICIER} \times ICIER \quad (17)$$

(16) Binding and unbinding between $ICI$ and $ER$

(17) Degradation of $ICIER$

$$\frac{dcyclinD1}{dt} = -kd_{cyclinD1} \times cyclinD1 \quad (18)$$

$$+k_{cyclinD1} + k_{cyclinD1E2ER} \times \frac{E2ER^{p_{cyclinD1E2ER_2}}}{p_{cyclinD1E2ER_1}^{p_{cyclinD1E2ER_2}} + E2ER^{p_{cyclinD1E2ER_2}}} \quad (19)$$

$$-kb_{cyclinD1cdk46} \times cyclinD1 \times cdk4 + kub_{cyclinD1cdk46} \times cyclinD1cdk4 \quad (20)$$

$$-kb_{cyclinD1cdk46} \times cyclinD1 \times cdk6 + kub_{cyclinD1cdk46} \times cyclinD1cdk6 \quad (21)$$

(18) Degradation of $cyclinD1$

(19) Basal translation of $cyclinD1$ and the increased by $E2ER$

(20) Binding and unbinding between $cyclinD1$ and $cdk4$

(21) Binding and unbinding between $cyclinD1$ and $cdk6$

$$\frac{drescdk6ICI}{dt} = k_{rescdk6ICI} \times \frac{ICI^{p_{rescdk6ICI_2}}}{p_{rescdk6ICI_1}^{p_{rescdk6ICI_2}} + ICI^{p_{rescdk6ICI_2}}} \quad (22)$$

$$-kd_{rescdk6ICI} \times \frac{rescdk6ICI^{p_{kdrescdk6ICI_2}}}{p_{kdrescdk6ICI_1}^{p_{kdrescdk6ICI_2}} + rescdk6ICI^{p_{kdrescdk6ICI_2}}} \quad (23)$$

$$-kd_{rescdk6ICI_{AZD}} \times rescdk6ICI \times sign(AZD) \quad (24)$$

(22) Increasing $rescdk6ICI$ by $ICI$

(23) Decreasing $rescdk6ICI$

(24) Decreasing $rescdk6ICI$ by $AZD$

$$\frac{dcdk4}{dt} = k_{cdk4} - kd_{cdk46} \times cdk4 \quad (25)$$

$$-kb_{cyclinD1cdk46} \times cyclinD1 \times cdk4 + kub_{cyclinD1cdk46} \times cyclinD1cdk4 \quad (26)$$

(25) Translation and degradation of $cdk4$

(26) Binding and unbinding between $cyclinD1$ and $cdk4$

$$\frac{dcdk6}{dt} = k_{cdk6} - kd_{cdk46} \times cdk6 \quad (27)$$

$$-kb_{cyclinD1cdk46} \times cyclinD1 \times cdk6 + kub_{cyclinD1cdk46} \times cyclinD1cdk6 \quad (28)$$

$$+k_{cdk6rescdk6ICI} \times \frac{rescdk6ICI^{p_{cdk6rescdk6ICI_2}}}{p_{cdk6rescdk6ICI_1}^{p_{cdk6rescdk6ICI_2}} + rescdk6ICI^{p_{cdk6rescdk6ICI_2}}} \quad (29)$$

(27) Translation and degradation of $cdk6$

(28) Binding and unbinding between $cyclinD1$ and $cdk6$

(29) Increasing $cdk6$ by $rescdk6ICI$

$$\frac{dcyclinD1cdk4}{dt} = -kd_{cyclinD1cdk46} \times cyclinD1cdk4 \quad (30)$$

$$+kb_{cyclinD1cdk46} \times cyclinD1 \times cdk4 - kub_{cyclinD1cdk46} \times cyclinD1cdk4 \quad (31)$$

$$-kb_{cyclinD1cdk46p21} \times cyclinD1cdk4 \times p21 + kub_{cyclinD1cdk46p21} \times cyclinD1cdk4p21 \quad (32)$$

$$-kb_{cyclinD1cdk46palbo} \times cyclinD1cdk4 \times palbo + kub_{cyclinD1cdk46palbo} \times cyclinD1cdk4palbo \quad (33)$$

(30) Degradation of $cyclinD1cdk4$

(31) Binding and unbinding between $cyclinD1$ and $cdk4$

(32) Binding and unbinding between $p21$ and $cyclinD1cdk4$

(33) Binding and unbinding between $palbo$ and $cyclinD1cdk4$

$$\frac{dcyclinD1cdk6}{dt} = -kd_{cyclinD1cdk46} \times cyclinD1cdk6 \quad (34)$$

$$+kb_{cyclinD1cdk46} \times cyclinD1 \times cdk6 - kub_{cyclinD1cdk46} \times cyclinD1cdk6 \quad (35)$$

$$-kb_{cyclinD1cdk46p21} \times cyclinD1cdk6 \times p21 + kub_{cyclinD1cdk46p21} \times cyclinD1cdk6p21 \quad (36)$$

$$-kb_{cyclinD1cdk46palbo} \times cyclinD1cdk6 \times palbo + kub_{cyclinD1cdk46palbo} \times cyclinD1cdk6dpalbo \quad (37)$$

(34) Degradation of $cyclinD1cdk6$

(35) Binding and unbinding between $cyclinD1$ and $cdk6$

(36) Binding and unbinding between $p21$ and $cyclinD1cdk6$

(37) Binding and unbinding between $palbo$ and $cyclinD1cdk6$

$$\frac{dcyclinD1cdk4p21}{dt} = -kd_{cdk46} \times cyclinD1cdk4p21 \quad (38)$$

$$+kb_{cyclinD1cdk46p21} \times cyclinD1cdk4 \times p21 - kub_{cyclinD1cdk46p21} \times cyclinD1cdk4p21 \quad (39)$$

$$-kb_{cyclinD1cdk46p21palbo} \times cyclinD1cdk4p21 \times palbo + kub_{cyclinD1cdk46p21palbo} \times cyclinD1cdk4p21palbo \quad (40)$$

(38) Degradation of $cyclinD1cdk4p21$

(39) Binding and unbinding between $p21$ and $cyclinD1cdk4$

(40) Binding and unbinding between $palbo$ and $cyclinD1cdk4p21$

$$\frac{dcyclinD1cdk6p21}{dt} = -kd_{cdk46} \times cyclinD1cdk6p21 \quad (41)$$

$$+kb_{cyclinD1cdk46p21} \times cyclinD1cdk6 \times p21 - kub_{cyclinD1cdk46p21} \times cyclinD1cdk6p21 \quad (42)$$

$$-kb_{cyclinD1cdk46p21palbo} \times cyclinD1cdk6p21 \times palbo + kub_{cyclinD1cdk46p21palbo} \times cyclinD1cdk6p21palbo \quad (43)$$

(41) Degradation of $cyclinD1cdk6p21$

(42) Binding and unbinding between $p21$ and $cyclinD1cdk6$

(43) Binding and unbinding between $palbo$ and $cyclinD1cdk6p21$

$$\frac{dcyclinD1cdk4palbo}{dt} = -kd_{cyclinD1cdk46} \times cyclinD1cdk4palbo \quad (44)$$

$$+kb_{cyclinD1cdk46palbo} \times cyclinD1cdk4 \times palbo - kub_{cyclinD1cdk46palbo} \times cyclinD1cdk4palbo \quad (45)$$

(44) Degradation of $cyclinD1cdk4palbo$

(45) Binding and unbinding between $palbo$ and $cyclinD1cdk4$

$$\frac{dcyclinD1cdk6palbo}{dt} = -kd_{cyclinD1cdk46} \times cyclinD1cdk6palbo \quad (46)$$

$$+kb_{cyclinD1cdk46palbo} \times cyclinD1cdk6 \times palbo - kub_{cyclinD1cdk46palbo} \times cyclinD1cdk6palbo \quad (47)$$

(46) Degradation of $cyclinD1cdk6palbo$

(47) Binding and unbinding between $palbo$ and $cyclinD1cdk6$

$$\frac{dcyclinD1cdk4p21palbo}{dt} = -kd_{cdk46} \times cyclinD1cdk4p21palbo \quad (48)$$

$$+kb_{cyclinD1cdk46p21palbo} \times cyclinD1cdk4p21 \times palbo - kub_{cyclinD1cdk46p21palbo} \times cyclinD1cdk4p21palbo \quad (49)$$

(48) Degradation of $cyclinD1cdk4p21palbo$

(49) Binding and unbinding between $palbo$ and $cyclinD1cdk4p21$

$$\frac{dcyclinD1cdk6p21palbo}{dt} = -kd_{cdk46} \times cyclinD1cdk6p21palbo \quad (50)$$

$$+kb_{cyclinD1cdk46p21palbo} \times cyclinD1cdk6p21 \times palbo - kub_{cyclinD1cdk46p21palbo} \times cyclinD1cdk6p21palbo \quad (51)$$

(50) Degradation of $cyclinD1cdk46p21palbo$

(51) Binding and unbinding between $palbo$ and $cyclinD1cdk46p21$

$$\frac{dcMyc}{dt} = k_{cMyc} - kd_{cMyc} \times cMyc \quad (52)$$

$$+k_{cMycE2ER} \times \frac{E2ER^{p_{cMycE2ER_2}}}{p_{cMycE2ER_1}^{p_{cMycE2ER_2}} + E2ER^{p_{cMycE2ER_2}}} + k_{cMycppRb} \times \frac{ppRb^{p_{cMycppRb_2}}}{p_{cMycppRb_1}^{p_{cMycppRb_2}} + ppRb^{p_{cMycppRb_2}}} \quad (53)$$

(52) Basal translation of $cMyc$ and degradation of $cMyc$

(53) Increase of $cMyc$ by $E2ER$ and $ppRb$

$$\frac{dp21}{dt} = k_{p21cMyc} \times \frac{p_{p21cMyc_1}^{p_{p21cMyc_2}}}{p_{p21cMyc_1}^{p_{p21cMyc_2}} + cMyc^{p_{p21cMyc_2}}} - kd_{p21} \times p21 \quad (54)$$

$$-kb_{cyclinD1cdk46p21} \times cyclinD1cdk4 \times p21 + kub_{cyclinD1cdk46p21} \times cyclinD1cdk4p21 \quad (55)$$

$$-kb_{cyclinD1cdk46p21} \times cyclinD1cdk6 \times p21 + kub_{cyclinD1cdk46p21} \times cyclinD1cdk6p21 \quad (56)$$

(54) Inhibited translation of $p21$ by $cMyc$ and degradation of $p21$

(55) Binding and unbinding between $p21$ and $cyclinD1cdk4$

(56) Binding and unbinding between $p21$ and $cyclinD1cdk6$

$$\frac{drescyclinE1palbo}{dt} = k_{rescyclinE1palbo} \times \frac{palbo^{p_{rescyclinE1palbo_2}}}{p_{rescyclinE1palbo_1}^{rescyclinE1palbo_2} + palbo^{p_{rescyclinE1palbo_2}}} \quad (57)$$

$$-kd_{rescyclinE1palbo} \times \frac{rescyclinE1palbo^{p_{kdrescyclinE1palbo_2}}}{p_{kdrescyclinE1palbo_1}^{p_{kdrescyclinE1palbo_2}} + rescyclinE1palbo^{p_{kdrescyclinE1palbo_2}}} \quad (58)$$

$$-kd_{rescyclinE1palbo_{AZD}} \times rescyclinE1palbo \times sign(AZD) \quad (59)$$

(57) Increasing $rescyclinE1palbo$ by $palbo$

(58) Decreasing $rescyclinE1palbo$

(59) Decreasing $rescyclinE1palbo$ by $AZD$

$$\frac{dcyclinE1}{dt} = -kd_{cyclinE1} \times cyclinE1 \quad (60)$$

$$+k_{cyclinE1} + k_{cyclinE1E2ER} \frac{E2ER^{p_{cyclinE1E2ER_2}}}{p_{cyclinE1E2ER_1}^{p_{cyclinE1E2ER_2}} + E2ER^{p_{cyclinE1E2ER_2}}} \quad (61)$$

$$+k_{cyclinE1rescyclinE1palbo} \times \frac{rescyclinE1palbo^{p_{cyclinE1rescyclinE1palbo_2}}}{p_{cyclinE1rescyclinE1palbo_1}^{p_{cyclinE1rescyclinE1palbo_2}} + rescyclinE1palbo^{p_{cyclinE1rescyclinE1palbo_2}}} \quad (62)$$

$$-kd_{cyclinE1_{AZD}} \times cyclinE1 \times sign(AZD) \quad (63)$$

(60) Degradation of $cyclinE1$

(61) Basal translation of $cyclinE1$ and increased by $E2ER$

(62) Increasing $cyclinE1$ by $rescyclinE1palbo$

(63) Decreasing $cyclinE1$ by $AZD$

$$\frac{dRb}{dt} = k_{Rb} - kd_{Rb} \times Rb \quad (64)$$

$$+k_{RbppRb} \times \frac{ppRb^{p_{RbppRb_2}}}{p_{RbppRb_1}^{p_{RbppRb_2}} + ppRb^{p_{RbppRb_2}}} \quad (65)$$

$$-k_{RbcyclinD1cdk4} \times \frac{cyclinD1cdk4^{p_{cyclinD1cdk4_2}}}{p_{cyclinD1cdk4_1}^{p_{cyclinD1cdk4_2}} + cyclinD1cdk4^{p_{cyclinD1cdk4_2}}} \times Rb \quad (66)$$

$$-k_{RbcyclinD1cdk6} \times \frac{cyclinD1cdk6^{p_{cyclinD1cdk6_2}}}{p_{cyclinD1cdk6_1}^{p_{cyclinD1cdk6_2}} + cyclinD1cdk6^{p_{cyclinD1cdk6_2}}} \times Rb \quad (67)$$

$$+k_{pRbdepho} \times pRb \quad (68)$$

(64) Basal translation and degradation of $Rb$

(65) Increasing $Rb$ by $ppRb$

(66) Phosphorylation of $Rb$ by $cyclinD1cdk4$

(67) Phosphorylation of $Rb$ by $cyclinD1cdk6$

(68) Dephosphorylation of $pRb$

$$\frac{dpRb}{dt} = -kd_{Rb} \times pRb \quad (69)$$

$$+k_{RbcyclinD1cdk4} \times \frac{cyclinD1cdk4^{p_{cyclinD1cdk4_2}}}{p_{cyclinD1cdk4_1}^{p_{cyclinD1cdk4_2}} + cyclinD1cdk4^{p_{cyclinD1cdk4_2}}} \times Rb \quad (70)$$

$$+k_{RbcyclinD1cdk6} \times \frac{cyclinD1cdk6^{p_{cyclinD1cdk6_2}}}{p_{cyclinD1cdk6_1}^{p_{cyclinD1cdk6_2}} + cyclinD1cdk6^{p_{cyclinD1cdk6_2}}} \times Rb \quad (71)$$

$$-k_{pRbdepho} \times pRb \quad (72)$$

$$-k_{pRbcyclinE1} \times \frac{cyclinE1^{p_{cyclinE1_2}}}{p_{cyclinE1_1}^{p_{cyclinE1_2}} + cyclinE1^{p_{cyclinE1_2}}} \times pRb \tag{73}$$

$$+k_{ppRbdepho} \times ppRb \tag{74}$$

(69) Degradation of $pRb$

(70) Phosphorylation of $Rb$ by $cyclinD1cdk4$

(71) Phosphorylation of $Rb$ by $cyclinD1cdk6$

(72) Dephosphorylation of $pRb$

(73) Phosphorylation of $pRb$ by $cyclinE$

(74) Dephosphorylation of $ppRb$

$$\frac{dppRb}{dt} = -kd_{ppRb} \times ppRb \tag{75}$$

$$+k_{pRbcyclinE1} \times \frac{cyclinE1^{p_{cyclinE1_2}}}{p_{cyclinE1_1}^{p_{cyclinE1_2}} + cyclinE1^{p_{cyclinE1_2}}} \times pRb \tag{76}$$

$$-k_{ppRbdepho} \times ppRb \tag{77}$$

(75) Degradation of $ppRb$

(76) Phosphorylation of $pRb$ by $cyclinE$

(77) Dephosphorylation of $ppRb$

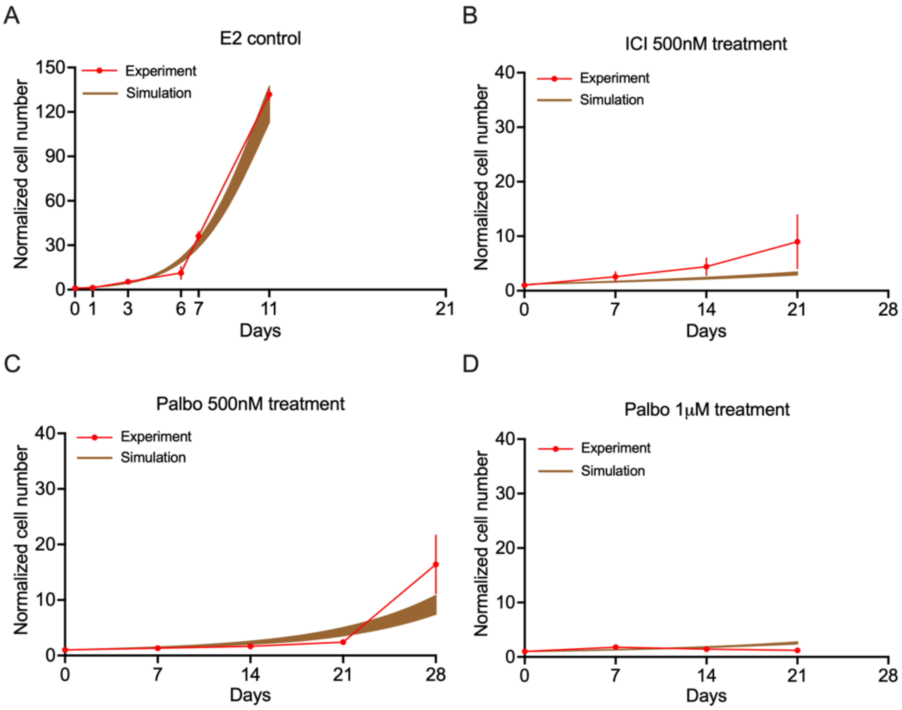

**Supplementary Figure 1. Mathematical model simulation compared to proliferation data for various treatments. (A)** No treatment (n = 3) for 11 days. The experimental data are shown in red and the simulation results are shown in brown (the shaded regions encompass the entire range of simulations within the cohort). **(B)** 500nM ICI treatment (n = 3) for 21 days. **(C)** 500nM Palbociclib treatment (n = 3) for 28 days. (**D)** 1μM Palbociclib treatment (n = 3) for 21 days. Palbo: palbociclib.

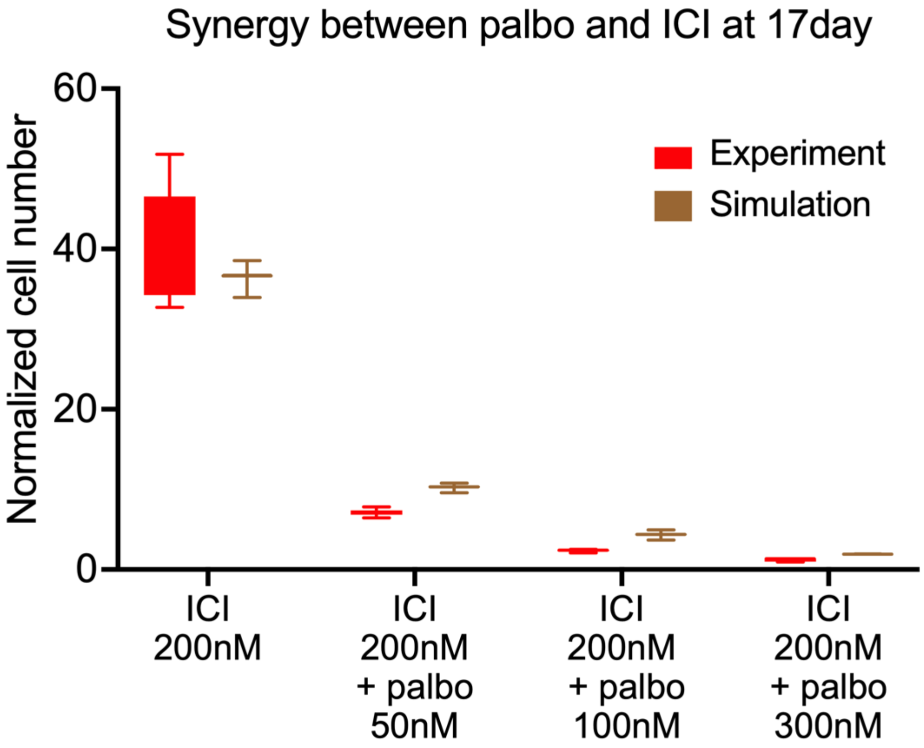


**Supplementary Figure 2. Boxplot of the model simulations and experimental verifications of normalized cell number showing the synergism between palbociclib and ICI.** The experimental results are shown in red. The simulation results are shown in brown from all cohort simulation results. The bottom and top lines on each box are the 25$^{th}$ and 75$^{th}$ percentiles, respectively. Palbo: palbociclib.
